# Population genomics of a bioluminescent symbiosis sheds light on symbiont transmission and specificity

**DOI:** 10.1101/736074

**Authors:** A.L. Gould, P.V. Dunlap

## Abstract

All organisms depend on symbiotic associations with bacteria for their success, yet the processes by which specific symbioses are established and persist remain largely undescribed. To examine the ecological mechanisms involved in maintaining symbiont specificity over host generations, we examined the population genomics of a binary symbiosis involving the coral reef cardinalfish *Siphamia tubifer* and the luminous bacterium *Photobacterium mandapamensis*. Using restriction site-associated sequencing (RAD-Seq) methods we demonstrate that the facultative symbiont of *S. tubifer* exhibits genetic structure at spatial scales of tens of kilometers in Okinawa, Japan in the absence of physical dispersal barriers and in contrast to the host fish. These results suggest the host’s behavioral ecology help structure symbiont populations at a reef site by symbiont enrichment, consequently fostering symbiont specificity. This approach also revealed several symbiont genes that were divergent between host populations including genes known to play a role in other host-bacteria associations.

## Introduction

In open marine environments, where there are few physical barriers to dispersal, microbes are expected to exist as well-mixed populations, supporting the classic panmictic view in microbial biogeography that “everything is everywhere, but the environment selects” ^1^. However, this view has been challenged in recent years, providing evidence of geographic structure in certain marine microbes^2–3^, although often over broad geographic scales and with coarse taxonomic resolution^4–7^. In contrast, there are few studies that have quantitatively examined the spatial scale of intra-specific patterns of distribution in marine bacteria with sufficient fine-scale genetic resolution to differentiate populations^8–10^. We define the population genomic structure of a symbiotically luminous marine bacterium over a relatively small geographic region and reveal the important role a host animal’s behavioral ecology can play symbiont transmission and in structuring populations of its microbial symbiont in a highly connected ocean environment.

The evolution of a symbiotic association requires that the specific partnership between host and symbiont remains stable over host generations. Such specificity is maintained in vertically transmitted symbioses via the intimate transfer of symbiont from parent to offspring and commonly leads to co-speciation^11–14^. In contrast, the evolution of horizontally transmitted symbioses requires both the selection of a particular symbiont from the environmental pool and the maintenance of that relationship over time. Therefore, a lower level of host-symbiont specificity is expected for horizontally acquired symbioses than that of their vertical counterparts. Nevertheless, many such associations are highly specific^15–17^, indicating that mechanisms such as environmental filtering and the ecology or physiology of the host ensure that a certain symbiont is acquired from the environment by each new generation.

Bioluminescent symbioses, especially those involving fish hosts, display a higher degree of specificity than what would be expected due to random symbiont acquisition from the marine environment^18–19^. How such specificity is achieved and maintained remains poorly understood, but the ecology of the host can play an important role^20–22^. Unfortunately, little is known of the ecology of most symbiotically luminous fishes, especially with respect to symbiont acquisition, a process that is challenging to study in nature due to the planktonic larval state of most fishes and their often deep or open water habitats^23^. The symbiotically luminous cardinalfish, *Siphamia tubifer*, inhabits shallow coral reefs in the Indo-Pacific and has known life history attributes^24^ and behavior^25^ that make it an ideal luminous fish species with which to investigate the host’s role in symbiont acquisition and in maintaining specificity of the symbiosis.

The bioluminescent symbiosis between *S. tubifer* and the luminous bacterium, *Photobacterium mandapamensis*, is a mutualistic association in which the symbiont is provisioned in an abdominal light organ by the fish in exchange for light production by the bacteria. The host, which remains quiescent among the long spines of sea urchins during the daytime^25–26^, uses the bacterially produced light to illuminate its ventral surface while foraging at night, possibly for countershading against the moonlight or for attracting prey. The host’s light organ is colonized by the luminous bacteria during larval development, although the exact timing in the wild remains unknown. In culture, *S. tubifer* larvae are not receptive to colonization by *P. mandapamensis* until at least eight days after having been released into the plankton^27^. Therefore, the direct transfer of symbiont from parent to offspring is not possible, and acquisition of the symbiont from the environment by aposymbiotic larvae occurs sometime prior to settlement, as the light organs of all juvenile *S. tubifer* collected from a reef have an established symbiont population. Several aspects of the initiation of the symbiosis remain undefined, including the timing and location of symbiont acquisition in the wild as well as the number of bacterial cells that initially colonize a light organ.

To gain a better understanding of symbiont acquisition for the *S. tubifer*-*P. mandapamensis* association we used a population genomics approach to quantify the within- and among- population genetic variability of the luminous bacteria from *S. tubifer* light organs collected from coral reefs in the Okinawa Islands, Japan. Specifically, we applied double digest restriction site-associated sequencing (ddRAD-Seq) methods to identify and analyze geographic patterns across hundreds of variant sites throughout the bacterial symbiont’s genome to test the hypothesis that the luminous symbionts of *S. tubifer* within a reef site are more genetically similar to each other than they are to the symbionts at other reefs. We also tested for evidence of population structure in the host fish and compared the patterns of genetic variability of the host to that of their symbiotic bacteria. We then analyzed the temporal stability of the light organ symbiont population at a single reef site over three consecutive years. This study provides insights into the process of symbiont acquisition for this highly specific, bioluminescent association, and we discuss the results in the context of the role of the host’s ecology in shaping the specificity of this horizontally transmitted symbiosis.

## Results

### Genetic admixture of the host fish

We carried out an analysis of the RAD-Seq data for a total of 282 *S. tubifer* specimens sampled from eleven locations that were separated by two to tens of kilometers (local scale) around Okinawa Island and up to 140 km (regional scale) between Okinawa Island and Kume Island. The host fish exhibited no evidence of genetic structure at either scale sampled in 2013 and 2014 (Figure 1) and all pairwise *F*_*ST*_ values were extremely low, ranging between 0.001 and 0.005 (SI Tables 1-3). These results confirm previous results that *S. tubifer* comprise one panmictic population in the Okinawa Islands likely due to larval dispersal^28^ during their month long pelagic stage^24^. Northward flow of the Kuroshio Current from the Philippine Islands promotes connectivity of marine populations in the region^29–30^, and a high frequency of typhoons and heavy predation on adult fish^24^, presumably all contribute to the observed genetic admixture in the region.

**Figure 1.**
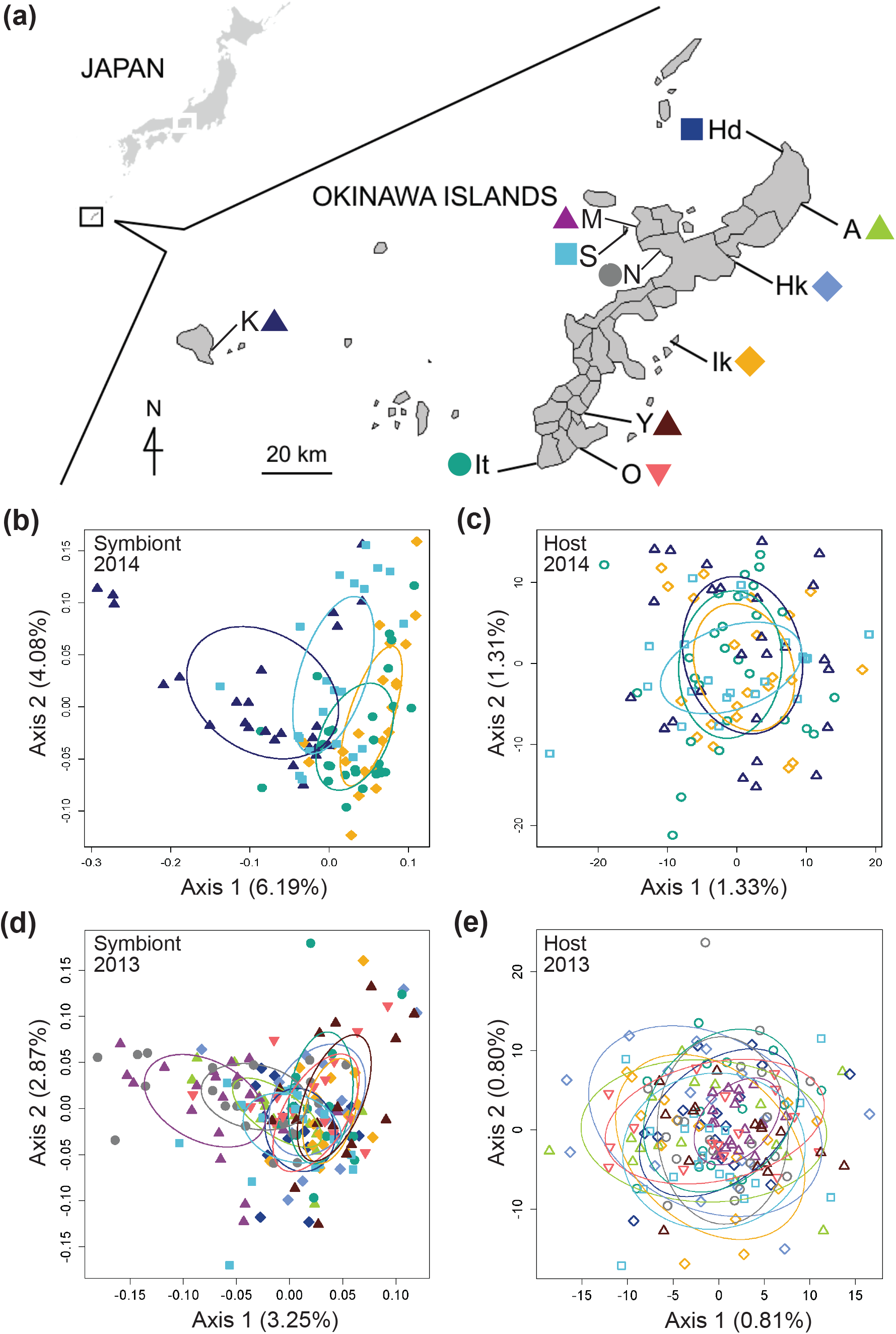
Genomic analysis of the light organ symbionts of *Siphamia tubifer* reveals structure in populations of symbiotic *Photobacterium mandapamensis* despite a lack of structure in the host fish. **(a)** Samples were collected from ten locations in 2013 and from four locations in 2014 in the Okinawa Islands, Japan. Principal coordinates analysis of genetic differentiation of **(b)** symbiotic *P. mandapamensis* across 534 loci analyzed from light organs sampled in 2014, **(c)** the corresponding *S. tubifer* hosts across 8,637 SNPs, **(d)** symbiotic *P. mandapamensis* across 465 loci analyzed from light organs sampled in 2013, and **(e)** the corresponding *S. tubifer* hosts across 8 637 SNPs. Points represent individuals along the first and second axes of genetic variation. The first 20 axes of variation for each analysis are depicted in SI Figure 5. Ellipses indicate standard deviation for symbiont populations sampled at each location.

### Population structure of the bacterial symbiont

Using a ddRAD-Seq approach, we identified a total of 601 bacterial loci for the light organ symbionts of *S. tubifer* across all 282 individuals with the restriction enzymes *Eco*RI and *Mse*I. These loci were distributed approximately evenly throughout the *P. mandapamensis* genome (Figure 2a,b) and thus, provide a representative snapshot of the symbiont genome. Ordination analyses of the filtered subset of these loci (n=534) from fish sampled in 2014 revealed a signature of genetic structure for populations of the symbiotic bacteria in contrast to the host fish (Figure 1), and was confirmed to be significant by PERMANOVA (Table 1). This structure was particularly evident when comparing symbiont populations between Okinawa Island and Kume Island, located approximately 140 km to the West (Figure 1b), which had significant differences in genetic distance to the light organ symbionts sampled from all three other locations (SI Table 4). However, loci analyzed (n=465) from light organs sampled in 2013 at the more local geographic scale of two to tens of kilometers suggest that the symbiotic bacteria are more genetically homogeneous around Okinawa Island (Table 1), although significant differences between populations in Okinawa were also detected (SI Table 5). Furthermore, no correlation in genetic distances was evident between the bacteria and their host fish for any of the three datasets (2013: r = −0.38, p = 1.00; 2014: r = −0.01, p = 0.53; Sesoko: r = −0.12, p = 0.89).

**Figure 2.**
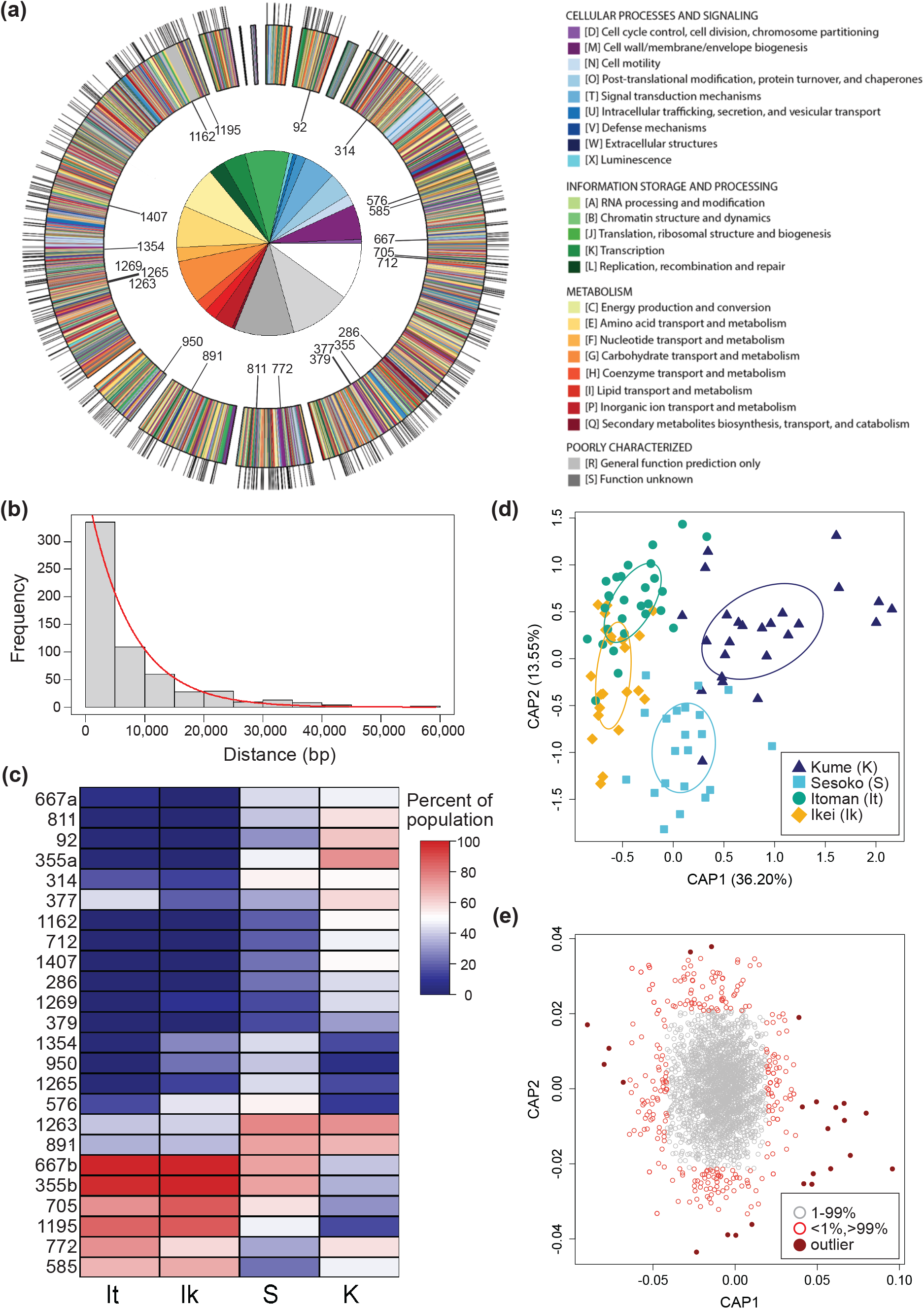
Restriction site-associated sequencing (RAD-Seq) yields a high density of genomic markers for the analysis of population structure and divergence of a symbiotic bacterium. **(a)** Annotated genome map of *Photobacterium mandapamensis*, the light organ symbiont of *Siphamia tubifer*. Protein-coding genes are color-coded according to which prokaryotic cluster of orthologous group (COG) they belong. Non-coding regions are white. Outer tic marks represent the locations of the 601 total loci analyzed for genomic differentiation between symbiont populations. The inner plot highlights the relative proportion of COG functions in which these loci are located. The 22 loci in which outlier haplotypes were identified as potential drivers of population differentiation are indicated on the inner circle of the map. **(b)** Distribution of the frequency of nucleotide distances between each pair of the 601 loci located in a scaffold of the *P. mandapamensis* genome. **(c)** Constrained analysis of principal coordinates (CAP) analysis of the 534 loci examined in 2014 **(d)** Corresponding CAP scores of the haplotypes analyzed across all 534 loci **(e)** Heatmap indicating the relative abundance within each population of the identified outlier haplotypes associated with each locus indicated.

**Table 1.** Permutational multivariate analyses of variance (PERMANOVA) of the genetic distances between *Siphamia tubifer* hosts and between their light organ symbionts. For each dataset the degrees of freedom (DF), sum of squares (SS), predicted F-value (F), R^2^ (R), and P-value are shown. For the 2014 and 2013 datasets, sampling location was used as the grouping factor, whereas sampling year was used for the Sesoko dataset. Significant P-values are in bold

| Dataset | Host |  |  |  |  | Symbiont |  |  |  |  |
| --- | --- | --- | --- | --- | --- | --- | --- | --- | --- | --- |
|  | DF | SS | F | R | P | DF | SS | F | R | P |
| **2014** | 3 | 15694 | 1.004 | 0.032 | 0.343 | 3 | 0.523 | 1.597 | 0.050 | **<0.001** |
| **2013** | 9 | 47367 | 1.008 | 0.054 | 0.131 | 9 | 1.099 | 1.249 | 0.064 | **<0.001** |
| **Sesoko** | 2 | 10718 | 1.027 | 0.038 | **0.033** | 2 | 0.225 | 1.092 | 0.043 | 0.093 |

### Loci correlated with symbiont population divergence

We identified putative loci driving the observed pattern of genetic divergence between populations of the symbiotic bacteria in 2014 and identified 24 outlier haplotypes from 22 loci. These haplotypes were highly correlated, either positively or negatively, with the primary axis of genetic variance differentiating the Kume Island and Okinawa Island populations (Figure 2c-e, SI Figure 4). All 22 loci were located in protein coding regions of the genome, and most of the variant effects examined for each of the outlier haplotypes were non-synonymous, including two that resulted in the addition of a stop codon (Table 2). Interestingly, some of these variants were in genes of known importance for host association in species of *Aliivibrio* and *Vibrio*, genera closely related to *Photobacterium*, including *hfq* (RNA-binding protein), *barA* (histidine kinase), and *gcvP* (glycine dehydrogenase), and could therefore, also be of importance for the bioluminescent association between *S. tubifer* and *P. mandapamensis*. Although the bacterial genes involved in the symbiosis have yet to be identified and little is known regarding the genetics of this association, except that homologs of non-*lux* genes involved in the bioluminescent symbiosis of *Aliivibrio fischeri* with the sepiolid squid, *Euprymna scolopes*, are absent in the *P. mandapamensis* genome^31^.

**Table 2.** Outlier haplotypes for *Photobacterium mandapamensis* identified as potential drivers of genetic divergence between symbiotic populations at Kume and Okinawa Islands, Japan in 2014. Listed are the locus ID assigned by *Stacks* (Catchen et al. 2011, 2013) corresponding to each haplotype, its variant type and effect as determined with *SnpEff*, and the putative gene product and category (COG) of the protein coding region in which each locus is located

| ID | Scaffold | Variant Type | Variant Effect | Gene | Gene product |
| --- | --- | --- | --- | --- | --- |
| 92 | 2 | missense | moderate | *hfq* | RNA chaperone Hfq |
| 286 | 4 | synonymous | LOW | PMSV_1336 | unknown function |
| 314 | 4 | synonymous | LOW | PMSV_145 | methyl-accepting chemotaxis protein |
| 355a | 4 | missense | moderate | PMSV_1452 | efflux RND transporter permease subunit |
| 355b |  | consensus | - |  |  |
| 377 | 4 | missense | moderate | *yeaG* | PrkA family serine protein kinase |
| 379 | 4 | missense | moderate | *yeaG* | PrkA family serine protein kinase |
| 576 | 4 | upstream | modifier | PMSV_554 | unknown function |
| 585 | 4 | missense | moderate | PMSV_564 | lipopolysaccharide assembly protein LapB |
| 667a | 4 | consensus | - | PMSV_772 | Hsp70 family protein |
| 667b |  | stop gained | high |  |  |
| 705 | 4 | missense | moderate | PMSV_851 | inosine/guanosine kinase |
| 712 | 4 | missense | moderate | VV12383 | DUF2786 domain-containing protein |
| 772 | 5 | missense | moderate | *hpt* | hypoxanthine phosphoribosyltransferase |
| 811 | 5 | missense | moderate | *barA* | two-component sensor histidine kinase BarA |
| 891 | 6 | missense | moderate | PMSV_3308 | Trk system potassium uptake protein |
| 950 | 7 | missense | moderate | PMSV_3736 | sigma-54-dependent Fis family transcriptional regulator |
| 1162 | 8 | missense | moderate | PMSV_2985 | unknown function |
| 1195 | 8 | consensus | - | PMSV_3012 | AsmA family protein |
| 1263 | 8 | missense | moderate | PMSV_2003 | alpha/beta fold hydrolase |
| 1265 | 8 | stop gained | high | PMSV_2003 | alpha/beta fold hydrolase |
| 1269 | 8 | synonymous | low | PMSV_2013 | cobalamin ABC transporter substrate-binding protein |
| 1354 | 8 | missense | moderate | PMSV_2243 | PTS fructose transporter subunit IIC |
| 1407 | 8 | missense | moderate | *gcvP* | aminomethyl-transferring glycine dehydrogenase |

### Temporal stability of symbiont populations

To determine the extent to which light organ symbiont populations were stable over time, we also analyzed the genetic variance of bacteria from fish light organs collected from one location (Sesoko, Figure 1a) in three consecutive years (2012, 2013, and 2014). In contrast to the genetic differentiation observed between sampling locations in 2014, populations of the symbiotic bacteria were not significantly different between years at this location (Figure 3, Table 1). This lack of variation suggests that the bacterial populations are somewhat stable at a reef over time. The host fish also exhibited little genetic differentiation between years (Figure 3), however a previous analysis of outlier loci from this dataset revealed a strong signature of temporal differentiation likely due to variable larval supply from different upstream sources^28^.

**Figure 3.**
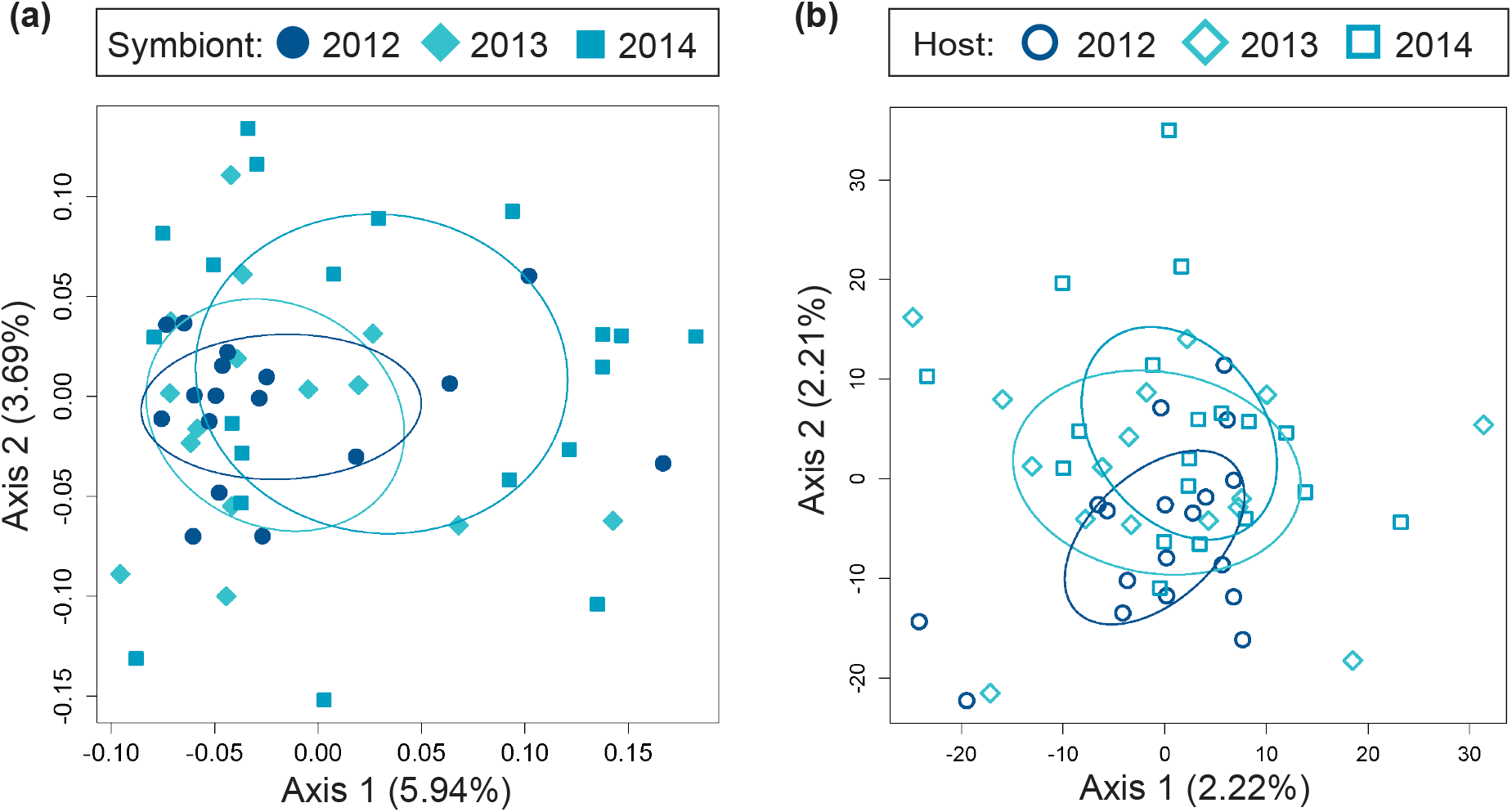
Temporal genomic analysis of the light organ symbionts of *Siphamia tubifer* reveals population stability at the same reef site over time. Light organs were sampled from reefs at Sesoko Island in Okinawa in the summer of 2012, 2013, and 2014. Principal coordinates analysis of genetic differentiation (**a**) of symbiotic *P. mandapamensis* across 552 loci and (**b**) of the corresponding *S. tubifer* hosts across 8,637 SNPs. Points represent individuals along the first and second axes of genetic variation. Ellipses indicate standard deviation for symbiont populations sampled each year.

### Within-population symbiont strain diversity

In addition to defining the genomic diversity of symbiotic *P. mandapamensis* populations between reefs, we also examined symbiont diversity within individual light organs. To estimate the number of genetically distinct bacterial strains comprising each light organ’s symbiont population, we determined the number of bacterial genotypes across all 601 loci within each host. However, one limitation of using RAD-Seq to analyze entire symbiont populations, is that haplotypes cannot be concatenated across loci. We therefore estimated the within light organ diversity to be the maximum number of haplotypes observed among all bacterial loci within a host. Using this approach, an average of six (± 1.6 SD) distinct bacterial genotypes were detected within a light organ (Figure 4), although this could be an underestimate of the total strain diversity present due to our inability to concatenate haplotypes. Individual hosts had anywhere between two and ten symbiont types present in their light organ, and no fish harbored a population comprised entirely of a single symbiont genotype (Figure 4). Furthermore, there was no correlation between fish length, a proxy for fish age^24^, and the minimum or maximum number of symbiont genotypes in a light organ (*F*_1,290_ = 1.477, *R*^2^_*Adj*_ = 0.00164, P = 0.225) (Figure 4), indicating the number of strains within a light organ does not increase over time. These results suggest that the population of bacteria within a light organ is likely founded by an average of six genetically distinct strains of *P. mandapamensis*, which then persist within the light organ throughout the host’s life.

**Figure 4.**
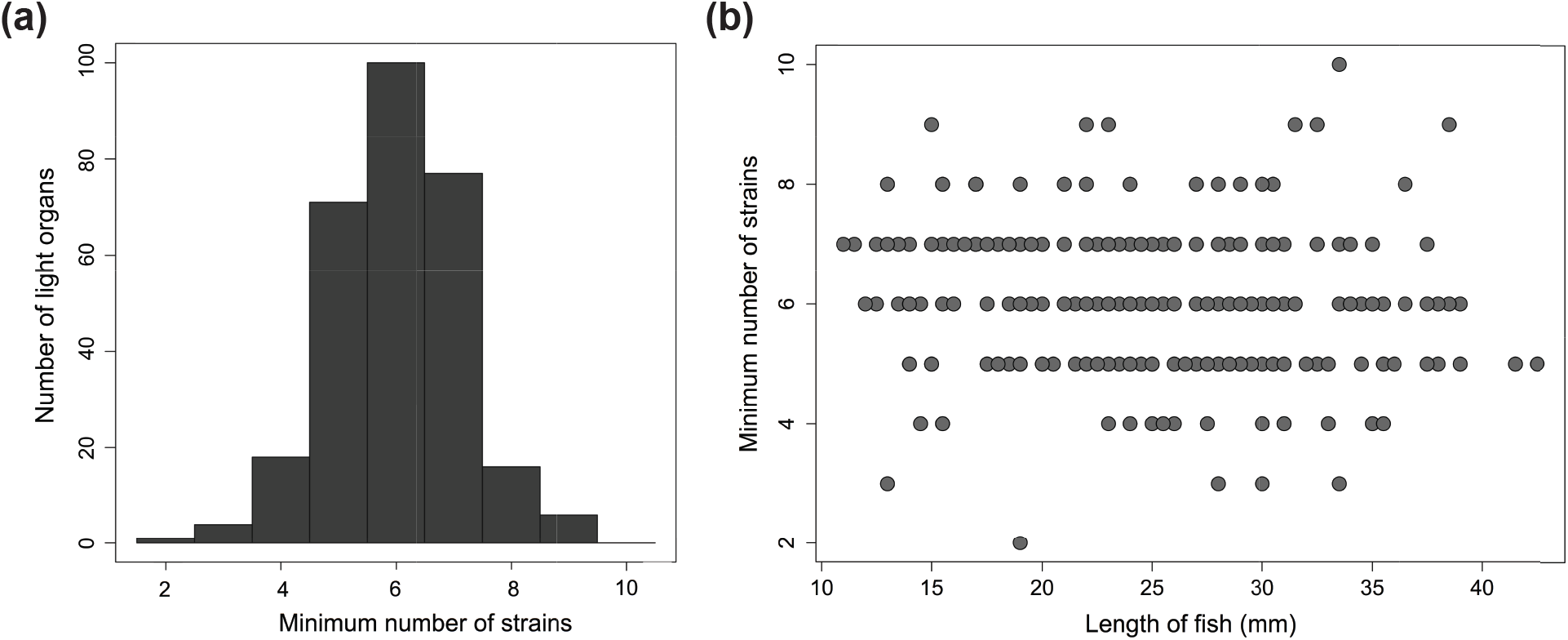
An average of six distinct *Photobacterium mandapamensis* strains are present within a light organ of *Siphamia tubifer*. **(a)** Frequency distribution of the minimum possible number of symbiotic *P. mandapamensis* genotypes present within a light organ as determined by the maximum number of haplotypes observed across all 601 symbiont loci analyzed. **(b)** The minimum number of distinct *P. mandapamensis* strains present within a light organ relative to the standard length of the host fish.

## Discussion

We obtained data for discrete, live populations of bacterial symbionts by extracting and sequencing genomic DNA directly from whole light organs of *S. tubifer*, a symbiotically luminous fish commonly found on coral reefs in the Okinawa Islands. This approach together with RAD-Seq methods allowed us to obtain deeply sampled, fine-scale genomic data capable of distinguishing populations of this bacterium from various locations sampled in the region, and in the absence of obvious barriers to dispersal. Although RAD-Seq methods have become increasingly popular in recent years^32–33^, they have not previously been used to examine the genomic structure of natural populations of bacteria nor have they been used to simultaneously examine populations of a host animal and its bacterial symbiont population. By applying these methods, we were able to identify potential genes of interest that are driving the patterns of genetic divergence between symbiont populations. Most notably, we discovered missense variations in *barA*, and *hfq* that were completely absent in the two of the four populations examined (Table 2, Figure 2e). The two-component *barA/uvrY* homologs *varS/varA* in *Vibrio cholerae* have been recognized as important for virulence^34–35^, quorum sensing^36^, and dissemination from a host to an aquatic environment^37^, and *varS* deletion strains of *V. cholerae* have colonization deficiencies in mice^35^. Similarly, the RNA-binding protein Hfq is essential for the virulence of *V. cholerae*, and strains lacking *hfq* also fail to colonize mice hosts^38^. Interestingly, *hfq* has also been identified as a negative regulator of bioluminescence in *V. harveyi*^39^. Additional variants of interest were observed in *gcvP*, which plays a role in the virulence of *V. cholerae* in *Drosophila* hosts^40^, and in *hpt*, which is located just upstream of the quorum-sensing regulating gene *luxR* in *V. harveyi*^41^. While the potential roles of these genes in the *S. tubifer*-*P. mandapamensis* symbiosis are unknown, our approach uncovered them as interesting targets for future studies of the underlying mechanisms regulating this bioluminescent association.

Despite the absence of physical barriers to dispersal and in contrast to the host fish, we discovered within-species level genetic differences between the luminous symbionts of *S. tubifer* sampled from different reefs in the Okinawa Islands. Furthermore, we determined that symbiont populations are seemingly stable at a reef between years. Environmental factors known to influence the biogeography of marine microbes include both temperature and salinity^2^. However, variation in these factors between sampling locations were unlikely to influence populations of symbiotic *P. mandapamensis* in this study due to the relatively small geographic area examined and shallow collection depths (<10 m). Therefore, we suggest the observed patterns of genetic structure between light organ symbionts from reefs in Okinawa and Kume Islands result from the behavioral ecology of the host.

The *S. tubifer* light organ is connected by a duct to the host’s intestine, and luminous bacterial cells are continually released via the duct from the light organ into the intestines and then the seawater with fecal waste^42^. The release of bacterial cells from the light organ presumably promotes the growth and light production of the symbiont population in the host while also enriching the surrounding seawater with symbiont cells^22,43–45^. Once settled as juveniles, *S. tubifer* exhibit fidelity to a home reef and typically return to the same urchin at that reef after foraging each night^25^. Therefore, the *P. mandapamensis* genotypes present in the light organs of resident fish at a reef are regularly released into the surrounding seawater and can become enriched in the local environment. Thus, the observed genetic structure of *P. mandapamensis* symbionts is likely a consequence of symbiont acquisition by larval fish occurring in the locally symbiont-enriched seawater near their settlement site. As a result, *S. tubifer* hosts residing at a particular reef, although not closely related, share more similar symbiont genotypes with each other than with fish from other reefs. This host-mediated symbiont population structure is consistent with the lack of temporal structure observed for symbiont populations sampled at the same reef over three years and has been observed for other marine symbioses^46^, including the bioluminescent symbiosis between *E. scolopes* and *A. fischeri*^47^.

In a highly connected ocean environment, where the classic view of microbial biogeography suggests that “everything is everywhere”^1^, we illustrate that fine-scale patterns of genomic population structure in a facultative bacterial symbiont can arise, even in a region dominated by strong ocean currents and in the absence of physical barriers to gene flow. These results provide new insight on the timing and location of symbiont acquisition by *S. tubifer* larvae in the wild, a process that is difficult to study for most horizontally acquired marine symbioses. We hypothesize that the distinct life history and behavioral ecology of *S. tubifer*^24–25^ results in the local enrichment of *P. mandapamensis* in the surrounding seawater, which consequently ensures that the next generation of host fish can successfully initiate its symbiosis with its species-specific luminous bacterial symbiont. This host facilitation presumably helps to foster the high degree of specificity observed for the partnership^19^ over time, and incidentally, can promote genetic divergence between symbiont populations. Future applications of RAD-Seq methods with other symbiotic associations have the potential to broaden our understanding of the influence of host animals on the biogeographic patterns of their microbial symbionts and the evolutionary processes that lead to host-symbiont integration and specificity.

## Materials and Methods

### Sampling, DNA extraction and library preparation

*Siphamia tubifer* specimens were collected over three years from locations in Okinawa, Japan (Table 3). Ten collection sites on Okinawa Island were sampled during June and July of 2013, and in June of 2014, samples were collected from an additional site on Kume Island as well as three sites on Okinawa Island that were previously sampled in 2013 (Figure 1a). One reef site (Sesoko, “S”) that was sampled in 2013 and 2014 was also sampled in July of 2012, providing a three-year dataset from that location. All collection sites were less than 10 m depth. Approximately 20 fish ranging in body lengths and associated with different sea urchins were collected at each sampling location. Fish were immediately euthanized upon collection by immersion in seawater containing a lethal dose (500 mg/L) of buffered tricaine methanesulfonate (MS-222) as approved by the University of Michigan Institutional Animal Care and Use Committee.

**Table 3.** Location, year, and number of *Siphamia tubifer* light organs sampled for this study in the Okinawa Islands, Japan.

| ID | Location | Latitude | Longitude | Year | N |
| --- | --- | --- | --- | --- | --- |
| S | Sesoko | 26.635409 | 127.865832 | 2012 | 16 |
|  |  |  |  | 2013 | 18 |
|  |  |  |  | 2014 | 21 |
| M | Motobu | 26.655806 | 127.880286 | 2013 | 20 |
| N | Nago | 26.603673 | 127.932404 | 2013 | 21 |
| Hd | Hedo | 26.848756 | 128.252513 | 2013 | 17 |
| It | Itoman | 26.095109 | 127.658478 | 2013 | 14 |
|  |  |  |  | 2014 | 27 |
| O | Ou | 26.127916 | 127.768981 | 2013 | 16 |
| Y | Yonabaru | 26.203032 | 127.771178 | 2013 | 16 |
| Ik | Ikei | 26.393535 | 127.988601 | 2013 | 15 |
|  |  |  |  | 2014 | 22 |
| Hk | Henoko | 26.534554 | 128.046181 | 2013 | 17 |
| A | Ada | 26.741936 | 128.321107 | 2013 | 16 |
| K | Kume | 26.351617 | 126.820149 | 2014 | 26 |

The whole, intact light organ, composed of fish tissue and containing the luminous bacterial symbiont population, was aseptically removed from each fish specimen and individually preserved in RNAlater®. Genomic DNA was extracted from each light organ using a QIAGEN DNeasy Blood and Tissue Kit following the manufacturer’s protocol, and double digest restriction site-associated sequencing (ddRAD-Seq) DNA libraries were built from the total DNA extracted from *S. tubifer* light organs, each individually barcoded with unique 10 bp DNA sequences for downstream identification. The protocol used to construct the ddRAD-Seq libraries followed a modified combination of previously described methods^48–49^, using the restriction enzymes *Mse*I and *Eco*RI, which were tested for their efficacy to produce a significant number of loci for the analysis of both the host fish and its symbiotic bacteria on an initial set of 50 light organs. Up to 50 DNA libraries were pooled per lane and sequenced at the Center for Applied Genomics, Toronto, ON, Canada, on the Illumina HiSeq2000 platform (San Diego, CA), generating 100 bp single-end sequence reads.

### Sequence processing and data analysis

Raw sequence reads were de-multiplexed and quality-filtered for a Phred score of 33 or higher with the *process_radtags* command in *Stacks* v1.35^50–51^. After the DNA sequence barcodes were removed, the remaining 90-bp sequence reads were aligned against the ∼4.5 Mb reference genome of *Photobacterium mandapamensis*^31^ using the *very_sensitive* command in *Bowtie2*^52^ v2.2.0 to separate the host fish sequences from those of its luminous symbiont. The unaligned sequences were additionally filtered against the genomes of both the marine bacterium *Vibrio campbelli* or *Escherichia coli* K12. All remaining unaligned sequences were classified as belonging to *S. tubifer*. Aligned *P. mandapamensis* sequences in *.SAM* file format were then additionally quality filtered using *SAMtools*^53^ v1.3, retaining only reads with a MAPQ score greater than 20. The quality-filtered, aligned *P. mandapamensis* sequences were then processed with the *ref_map* command in *Stacks*, requiring a minimum stack depth of three (-m 3). Individuals with fewer than 100,000 total sequence reads or a mean depth of coverage per stack less than 100x were discarded from the analysis as they were considered to be outliers with respect to evenness of coverage across the individual sequence libraries (SI Figure 1). The final dataset consisted of sequence reads from 282 light organs collected from 11 locations over three years in the Okinawa Islands (Table 3).

To minimize the effects of missing data, only bacterial loci present in at least 70% of all light organs were extracted from the entire catalog of loci produced by *Stacks* and used for the downstream analysis. Subsequently, loci not present in at least 50% of individuals in each population were removed from the dataset, resulting in an initial dataset of 601 RADtags (loci) across all individuals. Additional missing data filters were also tested but did not have a notable affect on the analysis (SI Figure 2), thus we present the results for the missing data filters described above. After this initial filtering step, individuals sampled in 2013, 2014, and from Sesoko (S) over three consecutive years (2012-14) were subsequently grouped into sub-datasets to analyze independently for genomic structure. To further minimize any potential effects of missing data within each dataset, both individuals and loci with more than 15% missing data were removed. The numbers of remaining loci analyzed were 465, 534, and 552 for the 2013, 2014, and Sesoko datasets, respectively. Within each locus, haplotypes with at least 6x coverage were then analyzed for patterns of genomic structure across light organ symbiont populations within each dataset. Additional haplotype depth filters (5×, 8×, and 10×) were also examined, but they also did not have a notable effect on the analysis (SI Figure 3). To visualize patterns of genetic variation in symbiont populations, principal coordinates analyses (PCoA) were performed on the Bray-Curtis dissimilarity matrices calculated from the presence or absence of these haplotypes across all light organs in each dataset. Significant differences in genetic differentiation were confirmed by a permutational multivariate analysis of variance (PERMANOVA) on the genetic dissimilarity matrices using the *adonis* function in the “vegan”^54^ package in R^55^ using sampling location (or year) as a factor. Pairwise *adonis* tests were then performed between all pairs of locations within the 2013 and 2014 datasets to determine which symbiont populations were significantly different from one another.

Patterns of genomic structure of the host fish were also analyzed and compared at the same scales as their light organ symbionts. *Siphamia tubifer* sequence reads were assembled *de novo* in *Stacks* and genotypes were assigned to each individual fish. Specifically, the first single nucleotide polymorphism (SNP) within each of the 8,637 loci present in 70% of individuals in each population with a minor allele frequency of 5% or greater were analyzed for each corresponding host fish. We selected these data filters based on previous RAD-Seq studies of other marine organisms^56–57^ in order to maximize the number of loci examined while minimizing the effects of missing data. Notably, the nonparametric analyses applied to these data tend to show little sensitivity to the minor allele filter^58^. We then performed principal coordinates analyses on the Euclidean genetic distances between individuals in each of the *S. tubifer* SNP datasets (2013, 2014, and Sesoko) using the “adegenet”^59^ package in R. Weir and Cockerham’s pairwise *F*_*ST*_ values^60^ between populations were calculated with 1,000 permutations in GenoDive, and corresponding p-values were adjusted for multiple comparisons using the Bonferroni method. To test for significant differences in genetic differentiation between sampling locations (or year) additional PERMANOVAs were performed on the genetic dissimilarity matrices as described above for the symbiotic bacteria. Mantel tests with 1,000 permutations were also performed on the dissimilarity matrices calculated for each dataset (2013, 2014, and Sesoko) to test for correlations between the genetic distances of the host fish to that of its light organ symbionts with the “vegan” package^54^ in R.

To identify loci driving the patterns of genetic divergence between light organ symbionts at Kume Island and Okinawa Island in 2014, a constrained analysis of principal coordinates (CAP) using the *capscale* function in the “vegan” package^54^ in R was performed on the 2014 *P. mandapamensis* genetic dissimilarity matrix. Haplotypes with the highest (>99%) and lowest (<1%) scores along first two axes of variation were identified and of these, 24 outlier haplotypes were further selected as potential outliers driving the observed genetic divergence between symbiont populations based on their large differences in CAP scores relative to all other haplotypes (SI Figure 4, Figure 2c-e). For each of these outlier haplotypes, the associated locus and gene function were identified, and their variant type and functional effect were determined using the program *SnpEff* ^61^.

## Acknowledgements

We thank Dr. Saki Harii and the staff of Sesoko Station, University of the Ryukyus for logistical assistance as well as Katherine Dougan and Maggie Grundler, University of Michigan (UM), for technical and field assistance. We gratefully acknowledge members of Dr. Lacey Knowles’ Lab (UM) for guidance and assistance with the RAD-Seq protocols and data analysis. We also thank Raquel Rivadeneira and the UM Genomic Diversity Lab for technical services and the Center for Applied Genomics for their sequencing services. Funding for this project was provided by National Science Foundation grant DEB-1405286.

