## Supplementary Information for "Population genomics of a bioluminescent symbiosis sheds light on symbiont transmission and specificity"

**SI Table 1.** Pairwise $F_{ST}$ values (bottom) and corresponding adjusted p-values (top) between populations of the host fish sampled at the local scale in 2013. P-values in bold remained significant after correcting for multiple comparisons (Bonferroni method)

| **2013** | M | S | Y | N | Ad | H | Hk | Ik | ItH | Ou |
|---|---|---|---|---|---|---|---|---|---|---|
| M | - | **0.045** | **0.045** | **0.045** | **0.045** | **0.045** | **0.045** | **0.045** | **0.045** | **0.045** |
| S | 0.003 | - | **0.045** | **0.045** | **0.045** | **0.045** | **0.045** | **0.045** | **0.045** | **0.045** |
| Y | 0.003 | 0.002 | - | **0.045** | **0.045** | **0.045** | **0.045** | **0.045** | 0.090 | **0.045** |
| N | 0.004 | 0.003 | 0.004 | - | **0.045** | **0.045** | **0.045** | **0.045** | **0.045** | **0.045** |
| Ad | 0.004 | 0.002 | 0.003 | 0.004 | - | **0.045** | **0.045** | **0.045** | **0.045** | **0.045** |
| H | 0.004 | 0.002 | 0.003 | 0.004 | 0.004 | - | **0.045** | **0.045** | **0.045** | 0.090 |
| Hk | 0.004 | 0.003 | 0.003 | 0.004 | 0.004 | 0.003 | - | **0.045** | **0.045** | **0.045** |
| Ik | 0.004 | 0.003 | 0.002 | 0.005 | 0.003 | 0.004 | 0.004 | - | **0.045** | 0.090 |
| ItH | 0.004 | 0.003 | 0.003 | 0.004 | 0.004 | 0.004 | 0.004 | 0.005 | - | 0.225 |
| Ou | 0.003 | 0.003 | 0.002 | 0.004 | 0.003 | 0.004 | 0.003 | 0.003 | 0.003 | - |

**SI Table 2.** Pairwise $F_{ST}$ values (bottom) and corresponding adjusted p-values (top) between populations of the host fish sampled at the regional scale in 2014. P-values in bold remained significant after correcting for multiple comparisons (Bonferroni method).

| **2014** | Ik | ItH | K | S |
|---|---|---|---|---|
| Ik | - | **0.024** | **0.006** | **0.006** |
| ItH | 0.001 | - | **0.012** | **0.006** |
| K | 0.002 | 0.001 | - | **0.006** |
| S | 0.002 | 0.003 | 0.003 | |

**SI Table 3.** Pairwise $F_{ST}$ values (bottom) and corresponding adjusted p-values (top) between the host fish populations sampled at Sesoko Island in Okinawa, Japan over three consecutive years. P-values in bold remained significant after correcting for multiple comparisons (Bonferroni method)

| **Sesoko** | 2012 | 2013 | 2014 |
|---|---|---|---|
| 2012 | - | **0.003** | **0.003** |
| 2013 | 0.004 | - | **0.003** |
| 2014 | 0.005 | 0.003 | - |

**SI Table 4.** Pairwise adonis PERMANOVAs of the genetic distances between the light organ symbionts of *Siphamia tubifer* between individuals sampled at all locations in 2014. For each model, the factors' degrees of freedom (DF), sum of squares (SS), predicted F-value (F), $R^2$ (R), and adjusted P-value (Holm-Bonferroni method) are shown. Significant values are in bold

| **Locations** | **DF** | **SS** | **F** | **R** | **P.adj** |
|---|---|---|---|---|---|
| **Ik - It** | 1 | 0.095 | 0.963 | 0.020 | 0.626 |
| **Ik - K** | 1 | 0.290 | 2.514 | 0.053 | **0.003** |
| **Ik - S** | 1 | 0.119 | 1.182 | 0.028 | 0.166 |
| **It - K** | 1 | 0.255 | 2.208 | 0.041 | **0.003** |
| **It - S** | 1 | 0.142 | 1.377 | 0.028 | **0.010** |
| **S - K** | 1 | 0.170 | 1.408 | 0.031 | **0.010** |

**SI Table 5.** Pairwise adonis PERMANOVAs of the genetic distances between the light organ symbionts of *Siphamia tubifer* between individuals sampled at all locations in 2013. For each model, the factors' degrees of freedom (DF), sum of squares (SS), predicted F-value (F), $R^2$ (R), and adjusted P-value (Holm-Bonferroni method) are shown. Significant values are in bold

| Locations | DF | SS | F | R | P.adj |
|---|---|---|---|---|---|
| **A - H** | 1 | 0.103 | 1.097 | 0.034 | 0.679 |
| **A - Hk** | 1 | 0.106 | 1.110 | 0.035 | 0.679 |
| **A - Ik** | 1 | 0.112 | 1.215 | 0.040 | 0.225 |
| **A - It** | 1 | 0.102 | 1.085 | 0.037 | 0.679 |
| **A - M** | 1 | 0.150 | 1.387 | 0.039 | **0.040** |
| **A - N** | 1 | 0.137 | 1.328 | 0.037 | **0.018** |
| **A - O** | 1 | 0.121 | 1.287 | 0.041 | 0.119 |
| **A - S** | 1 | 0.101 | 1.067 | 0.032 | 0.679 |
| **A - Y** | 1 | 0.116 | 1.294 | 0.041 | 0.210 |
| **H - Hk** | 1 | 0.106 | 1.117 | 0.034 | 0.653 |
| **H - Ik** | 1 | 0.090 | 0.988 | 0.032 | 1.000 |
| **H - It** | 1 | 0.097 | 1.042 | 0.035 | 1.000 |
| **H - M** | 1 | 0.173 | 1.617 | 0.044 | **0.009** |
| **H - N** | 1 | 0.138 | 1.355 | 0.036 | **0.009** |
| **H - O** | 1 | 0.096 | 1.029 | 0.032 | 1.000 |
| **H - S** | 1 | 0.092 | 0.980 | 0.029 | 1.000 |
| **H - Y** | 1 | 0.107 | 1.212 | 0.038 | 0.245 |
| **Hk - Ik** | 1 | 0.087 | 0.933 | 0.030 | 1.000 |
| **Hk - It** | 1 | 0.082 | 0.874 | 0.029 | 1.000 |
| **Hk - M** | 1 | 0.196 | 1.809 | 0.049 | **0.009** |
| **Hk - N** | 1 | 0.137 | 1.328 | 0.036 | **0.016** |
| **Hk - O** | 1 | 0.091 | 0.967 | 0.030 | 1.000 |
| **Hk - S** | 1 | 0.111 | 1.161 | 0.034 | 0.420 |

| | | | | | |
|---|---|---|---|---|---|
| **Hk - Y** | 1 | 0.083 | 0.920 | 0.029 | 1.000 |
| **Ik - It** | 1 | 0.077 | 0.846 | 0.030 | 1.000 |
| **Ik - M** | 1 | 0.195 | 1.837 | 0.053 | **0.009** |
| **Ik - N** | 1 | 0.139 | 1.378 | 0.039 | **0.009** |
| **Ik - O** | 1 | 0.089 | 0.980 | 0.033 | 1.000 |
| **Ik - S** | 1 | 0.100 | 1.084 | 0.034 | 1.000 |
| **Ik - Y** | 1 | 0.068 | 0.783 | 0.026 | 1.000 |
| **It - M** | 1 | 0.171 | 1.581 | 0.047 | **0.009** |
| **It - N** | 1 | 0.124 | 1.212 | 0.035 | 0.128 |
| **It - O** | 1 | 0.071 | 0.768 | 0.027 | 1.000 |
| **It - S** | 1 | 0.081 | 0.860 | 0.028 | 1.000 |
| **It - Y** | 1 | 0.067 | 0.764 | 0.027 | 1.000 |
| **M - N** | 1 | 0.129 | 1.137 | 0.028 | 0.076 |
| **M - O** | 1 | 0.171 | 1.593 | 0.045 | **0.009** |
| **M - S** | 1 | 0.156 | 1.452 | 0.039 | **0.009** |
| **M - Y** | 1 | 0.198 | 1.915 | 0.053 | **0.009** |
| **N - O** | 1 | 0.132 | 1.295 | 0.036 | **0.005** |
| **N - S** | 1 | 0.133 | 1.293 | 0.034 | **0.003** |
| **N - Y** | 1 | 0.146 | 1.485 | 0.041 | **0.002** |
| **O - S** | 1 | 0.100 | 1.057 | 0.032 | 1.000 |
| **O - Y** | 1 | 0.090 | 1.023 | 0.033 | 1.000 |
| **S - Y** | 1 | 0.104 | 1.154 | 0.035 | 0.470 |

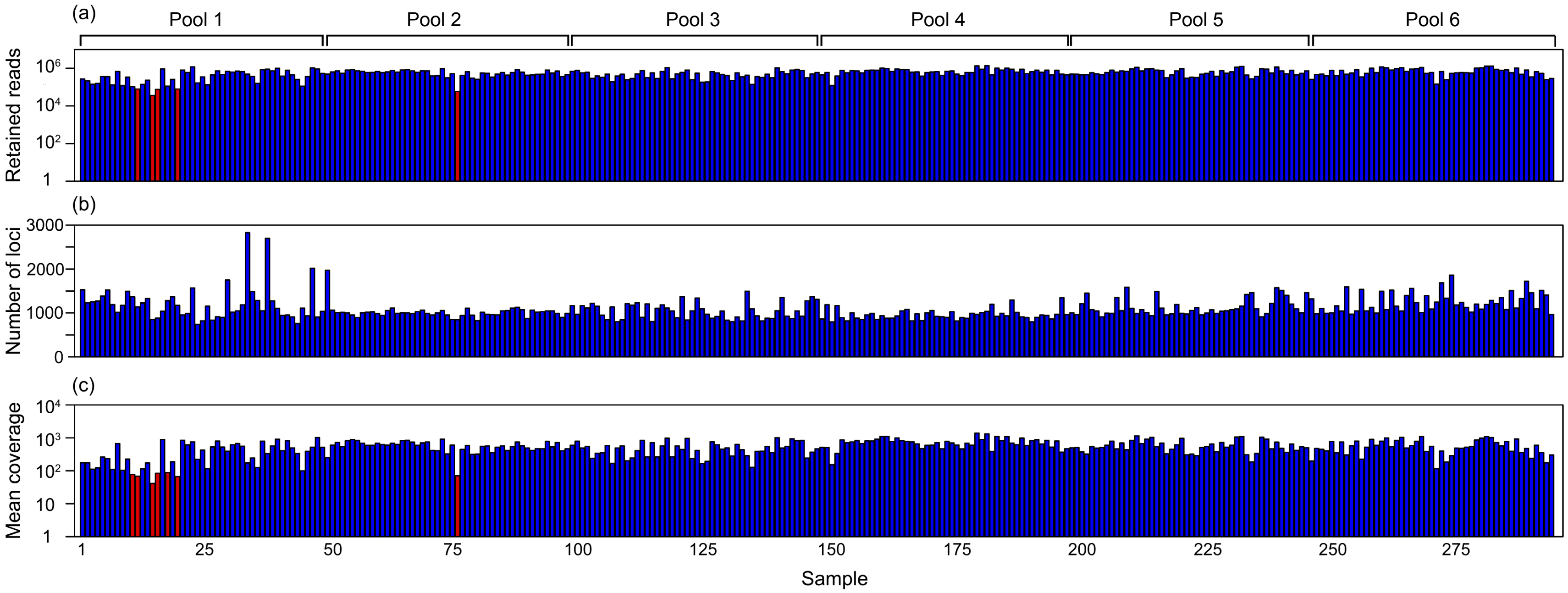


**SI Figure 1. The double digest restriction site associated sequencing (ddRAD-Seq) libraries produced high quality *Photobacterium mandapamensis* sequence reads from nearly all of the *Siphamia tubifer* light organs sampled**. Each bar represents symbiont sequences reads from an individual light organ library. **(a)** The total number of *P. mandapamensis* reads per light organ retained after quality filtering in *Stacks* (Catchen et al. 2011, 2013). **(b)** The total number of loci identified across the symbiont population per light organ using the *ref_map* command in *Stacks*. **(c)** The mean coverage of each locus per light organ. Bars highlighted in red are individuals that were removed from the analysis due to poor quality (too few reads or low depth of coverage).

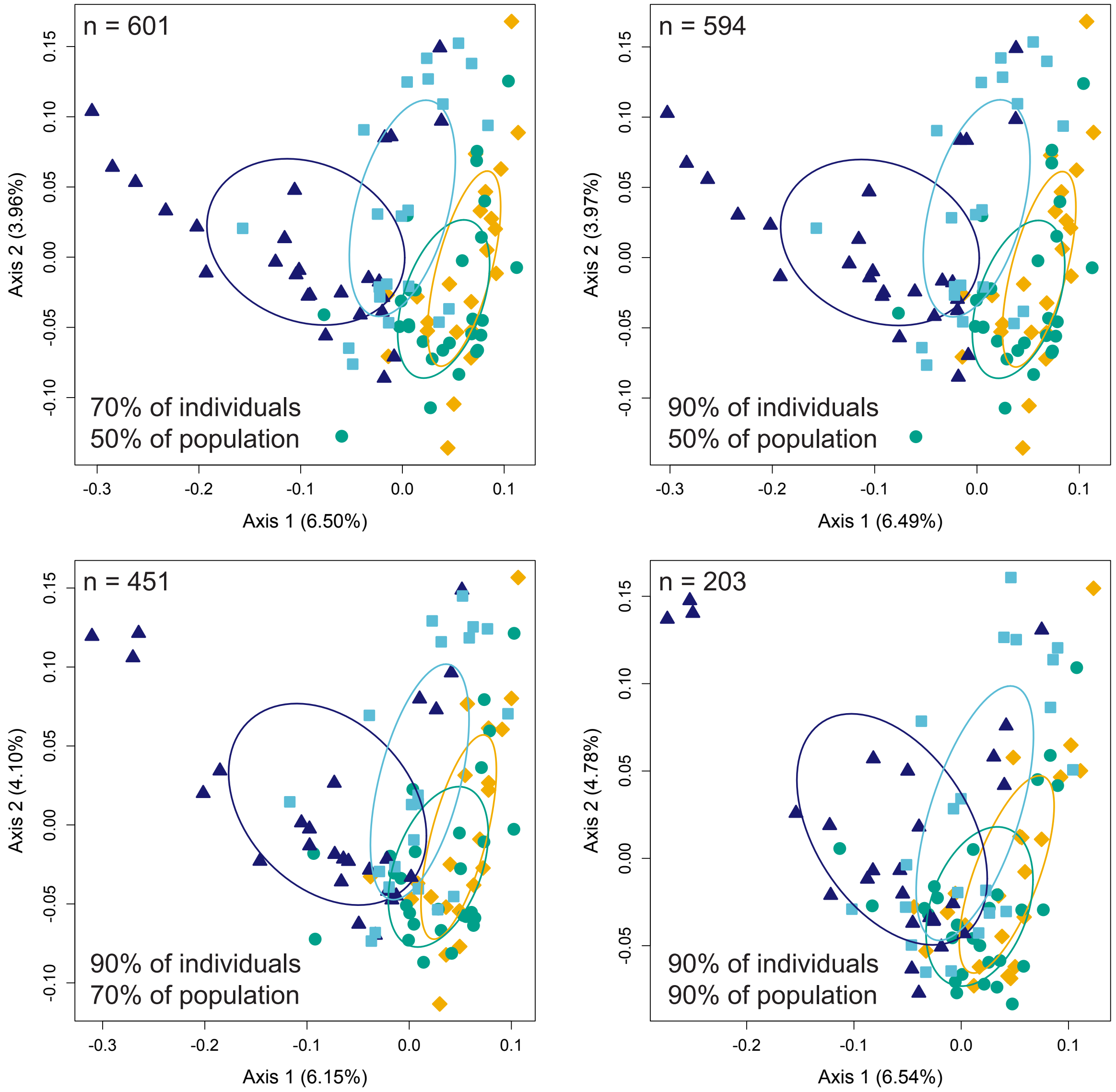


**SI Figure 2. Applying different missing data filters to the principal coordinate analyses of the genetic dissimilarities between the light organ symbionts of *Siphamia tubifer* have little effect on the patterns of genetic differentiation between sampling locations in 2014.** The total percent of all individuals sampled and the percent of individuals per population required for a locus to be included in the analysis are indicated in the bottom left corner of each plot. The number of loci analyzed is indicated in the top left corner

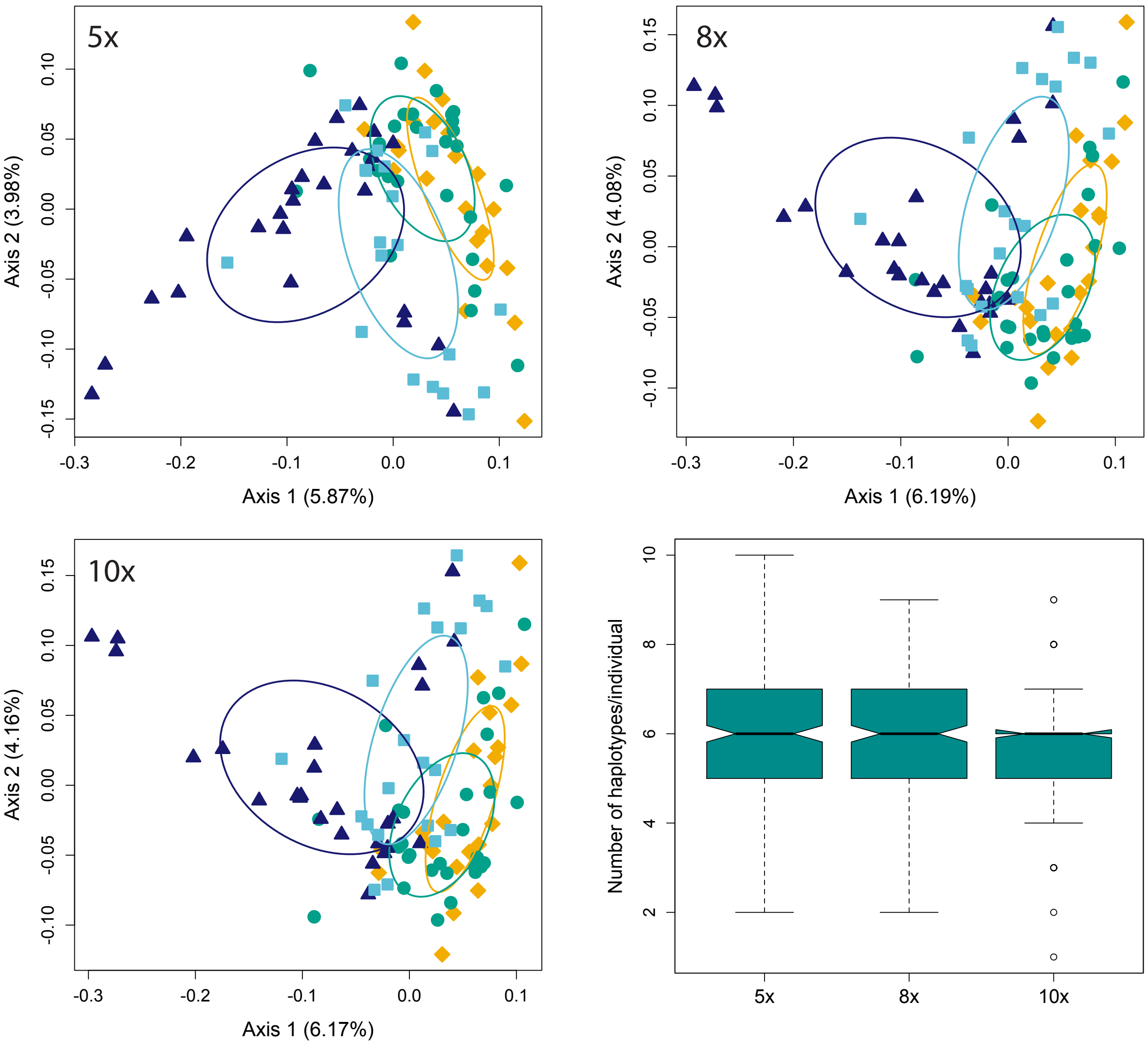


**SI Figure 3. Applying different haplotype depth filters to the principal coordinate analyses of the genetic dissimilarities between the light organ symbionts of *Siphamia tubifer* have little effect on the patterns of genetic differentiation between sampling locations in 2014.** The sequence depth required for a haplotype at any given locus to be included in the analysis is indicated in the top left corner of each plot. The distribution of the number of haplotypes per light organ, estimated as the maximum number of haplotypes present across all loci for an individual, when applying the depth filters indicated to call haplotypes

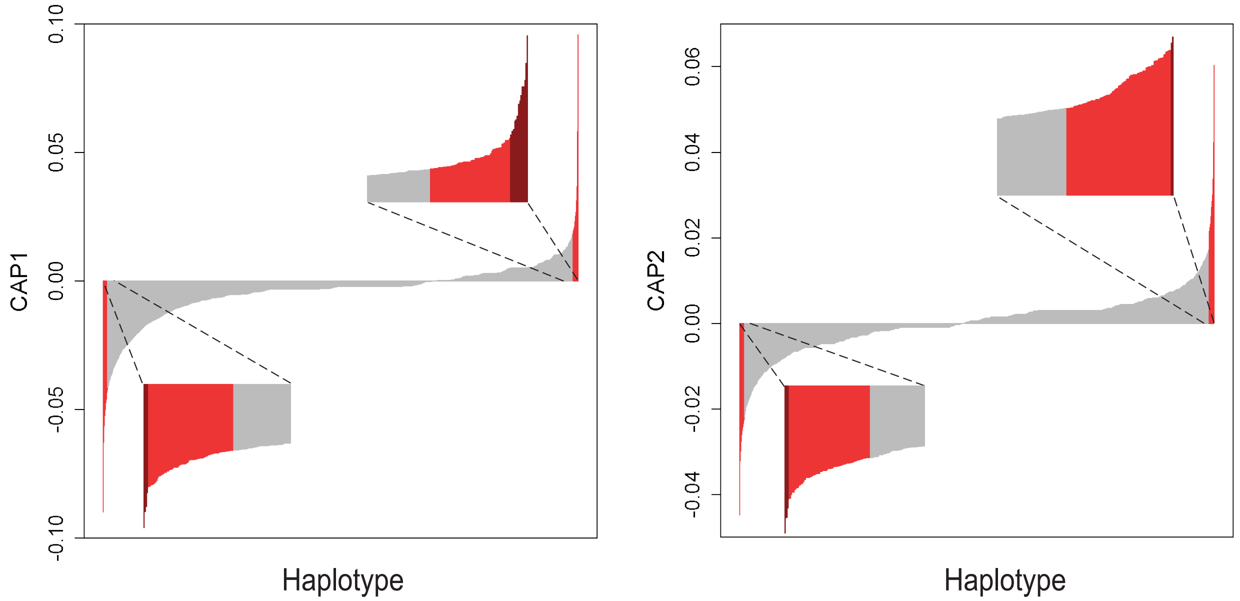


**SI Figure 4. The ranked distribution of the corresponding CAP1 (left) and CAP2 (right) scores for each haplotype in the constrained analysis of principal coordinates (CAP) analysis of the 551 loci examined in 2014.** Haplotypes with the highest and lowest 1% of scores are highlighted in red. Haplotypes analyzed as outliers are highlighted in dark red

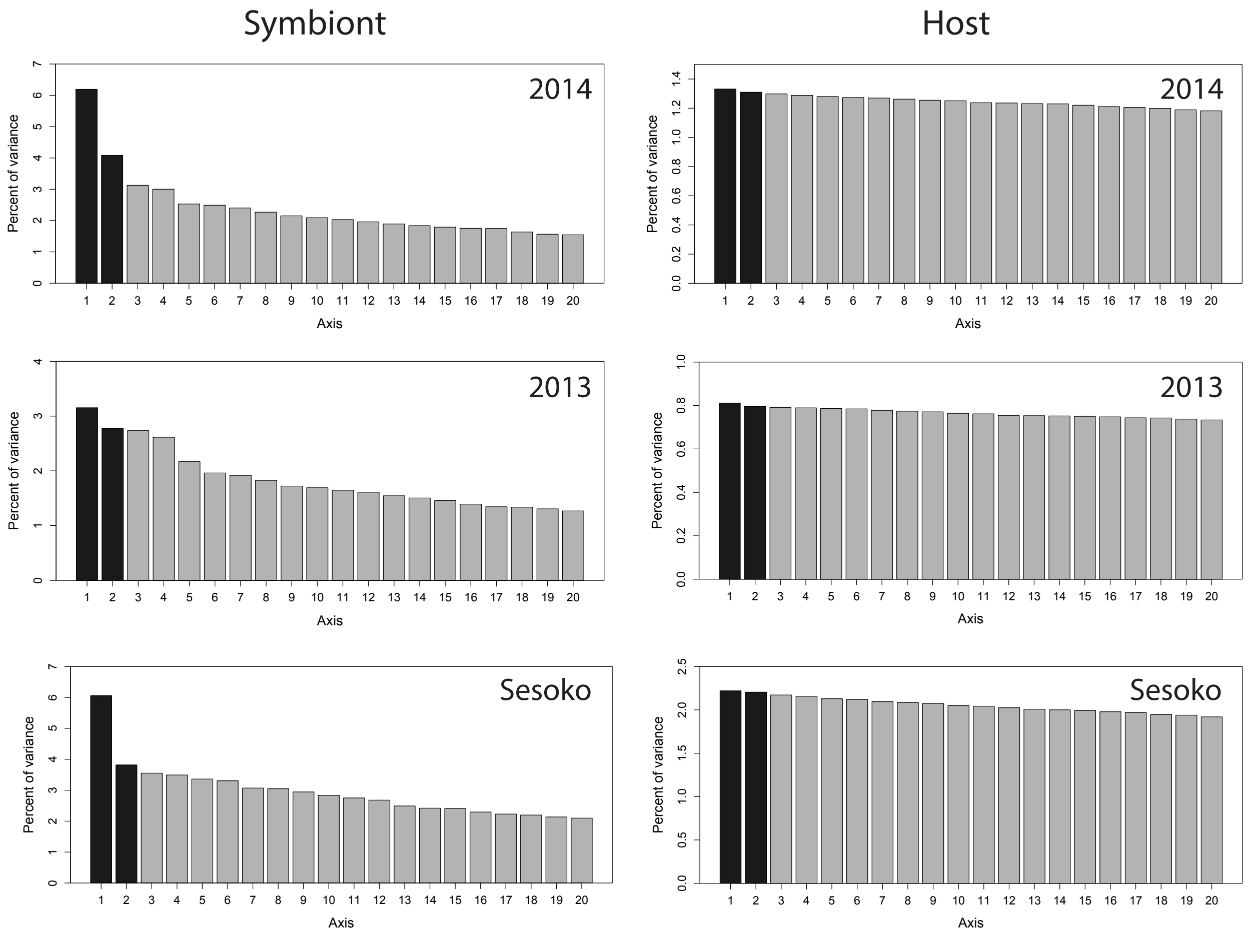


**SI Figure 5. The first 20 axes of variation for the principal coordinates analyses of genetic variation of the light organ symbionts of *Siphamia tubifer* (left column, "Bacteria") and the host fish (right column, "Fish") for the datasets indicated in the top right corner of each plot**
